# Personalized computational hemodynamic analysis in transcatheter aortic valve: investigation of long-term degeneration

**DOI:** 10.1101/2025.10.09.681392

**Authors:** Luca Crugnola, Chiara Catalano, Laura Fusini, Salvatore Pasta, Gianluca Pontone, Christian Vergara

## Abstract

Introduced as an alternative to open-heart surgery for elderly patients, Transcatheter Aortic Valve Implantation (TAVI) has recently been extended to younger patients due to comparable performance with the gold-standard. However, the long-term durability of the bio-prosthetic TAVI valves is limited by Structural Valve Deterioration (SVD), an inevitable degenerative process whose pathogenesis is still unclear. In this study, we aim to computationally investigate a possible relation between aortic hemodynamics and SVD development. To this aim, we collect data from twelve patients with and without SVD at long-term follow-up exams. Starting from pre-operative clinical images, we build early post-operative virtual scenarios and we perform Computational Fluid Dynamics simulations by prescribing a personalized flow rate based on Echo Doppler data. In order to identify a premature onset of SVD, we propose three computational hemodynamic indices: Wall Damage Index (*WDI*), Leaflet Delamination Index (*LDI*), and Leaflet Permeability Index (*LPI*). Additionally, to each index we associate a score and, using the Wilcoxon rank-sum test, we find that each score individually shows a statistically greater median value in the SVD sub-population (*WDI*: *p* = 0.008, *LDI*: *p* = 0.001, *LPI*: *p* = 0.020). Finally, we define a synthetic scoring system that clearly separates between SVD and non-SVD patients. Our results suggest that aortic hemodynamics may drive a premature onset of SVD, and the synthetic score could potentially assist clinicians in a patient-specific planning of follow-up exams to closely monitor those patients at high SVD risk.

## 1. Introduction

During the last two decades Transcatheter Aortic Valve Implantation (TAVI) has been established as a minimally invasive treatment of severe aortic stenosis in old patients at high-risk for open-heart surgery [1, 2]. Thanks to the positive outcomes of recent clinical trials [3, 4], nowadays TAVI represents the therapy of choice also in many lower-risk, younger patients [5]. In this context, it is particularly important to assess the long-term durability of the bio-prosthetic valves used for TAVI. Nonetheless, the relatively young age of this procedure, the limited life expectancy of its recipients, and the dispersion of patients after the intervention result in a lack of long-term follow-up data which makes it difficult to properly investigate the durability of TAVI valves.

The main limiting factor to the durability of bio-prosthetic valves is Structural Valve Deterioration (SVD), an intrinsic degenerative process manifested as permanent changes in the prosthesis, such as leaflet calcification or destruction of connective tissue, that ultimately leads to the failure of the implanted valves [5, 6]. However, the pathogenesis of the SVD process is still incompletely understood and multiple mechanisms are thought to be involved [7]. Specifically, an elevated host cell infiltration was observed in explanted bio-prosthetic valves with SVD [8, 9].

In the TAVI context, clinical studies proposed several SVD risk factors that can be divided into valve-related and patient-related factors [10, 11, 12]. Some of these factors are associated to a high cardiac output and a small valvular orifice area. Accordingly, Ochi et al. hypothesized that an accelerated flow across the valve is a shared key component in the pathophysiology of SVD [13]. This suggests an influence of hemodynamics on the development of SVD.

To better investigate this point, in this paper we assess the role of hemodynamics in SVD by means of computational modeling, which represents a powerful tool to obtain a detailed quantitative description of blood-dynamics after TAVI. Within the TAVI framework, computational models have been extensively employed to simulate the deployment of the bio-prosthetic valve [14, 15] and post-TAVI blood-dynamics [16, 17], focusing on post-procedural complications such as paravalvular leakage [18, 19], conduction abnormalities [20, 21], and coronary obstruction [22]. Additionally, the durability of bio-prosthetic valves has been associated to the distribution of mechanical stresses in different device designs using computational mechanical models [23, 24, 25]. Moreover, a recent work experimentally showed that SVD is associated with mineral precipitation and cellular infiltration in the bio-prosthetic leaflets and the authors proposed computational hemodynamic indices that correlate with their findings in idealized geometries [26]. However, to the best of our knowledge, the only computational hemodynamic studies investigating the relationship between aortic hemodynamics and SVD using patient-specific clinical data have been carried out in our previous studies [27, 28].

In this work we expand the analyses of [29, 28] by further refining and personalizing the Computational Fluid Dynamics (CFD) model. In particular, the advancements of this paper rely on the introduction of a wire-frame description of the bio-prosthetic valve’s stent, the modeling of the opening/closure dynamics of the valve’s leaflets, the use of realistic TAVI leaflets geometry, and the flow rate conditions derived by Echo Doppler data.

The aims of this retrospective computational study are to:

1. propose a framework to numerically simulate highly personalized post-TAVI hemodynamics starting from pre-operative clinical images and early post-operative Echo Doppler data;
2. identify new hemodynamic indices in early post-TAVI virtual scenarios, i.e. when the implanted valve has not degenerated yet, that correlate with a premature onset of SVD, detected at 5-10 years follow-up exam.

## 2. Materials and methods

### 2.1. Clinical data

In this study we leverage clinical data related to twelve patients who underwent the TAVI procedure in Centro Cardiologico Monzino between 2008 and 2012. In each patient it was implanted an Edwards SAPIEN valve of either first or second generation with a nominal size of 23 *mm* in external diameter. These are trileaflet bio-prosthetic valves made of bovine pericardium mounted on a balloon-expandable cylindrical stent with an inner fabric skirt on its ventricular side [30] (see Figure 1 left).

**Figure 1:**
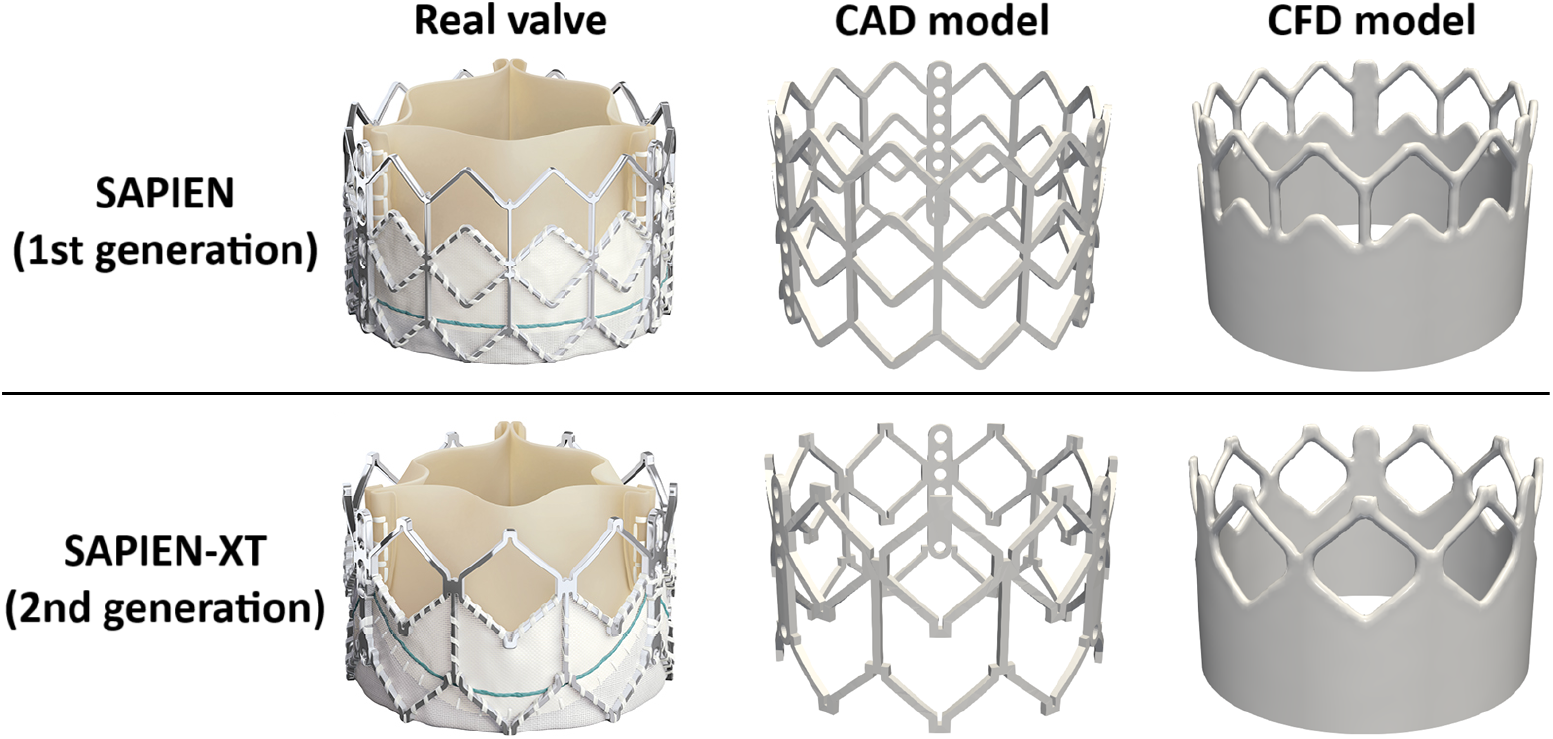
For each SAPIEN valve considered in this study, we report: an image of the real valve (left), the CAD model (center) and the stent model used in the CFD simulations (right). The “Real valve” images are taken from Rheude, Blumenstein, Möllmann, Husser under CC-BY-NC license, ©Edwards Lifensciences Corporation.

In this paper we adopt a definition of SVD as “stage 3 SVD” according to [31]. In particular, six patients with stage 3 SVD at long-term follow-up were identified and six patients without SVD at same exam were randomly extracted from the same cohort and matched for baseline characteristics.

For each patient we have at disposal the following data that we use as input in our model:

1. Pre-TAVI: Computed Tomography Angiography (CTA) scan. This is used in our model to reconstruct the patient-specific aortic geometries (see Section 2.2). See [11, 28] for details on the CTA acquisition;
2. Early post-TAVI: Pulsed Wave (PW) Doppler Transthoracic Echocardiography (TTE) data obtained between two days and seven months after the implantation. We exploit these data to derive a personalized flow rate profile that we impose as boundary condition in the numerical simulations (see Sections 2.3-2.4). See [11, 28] for details on the Doppler TTE acquisition;
3. Long-term follow-up: Doppler TTE data obtained 5-10 years after TAVI. These data are used to identify which patients developed SVD. The acquisition details are the same as in the previous point.

Moreover, for eleven out of twelve patients we have at disposal early post-TAVI maximum blood velocity measurements through the bio-prosthetic aortic valve taken from Continuous Wave (CW) Doppler TTE data that we use to validate the numerical hemodynamic results.

The study was approved from IRB of Centro Cardiologico Monzino and registered with number R1264/20.

### 2.2. Post-operative virtual scenarios

We aim to perform a personalized computational analysis of post-TAVI hemodynamics exploiting the available clinical data (Section 2.1), so that we build early post-TAVI virtual scenarios starting from pre-TAVI CTA scans. This procedure is extensively based on the use of the *Vascular Modeling Toolkit* (*VMTK*) [32] and *Paraview* open-source software and was already presented in [29, 28]. In the following we give a brief overview of the procedure, detailing the advancements introduced in this study.

Pre-operative patient-specific geometries of the aortic root, the ascending aorta and the aortic valve’s calcium deposits are semi-automatically reconstructed. The reconstructed aortic root geometry is then possibly deformed to be able to host the bio-prosthetic valve model, coherently to what happens during the balloon inflation step of the TAVI procedure. A cylindrical model of the SAPIEN valve’s stent is oriented according to the aortic root centerline, centered in the aortic annulus barycenter and virtually inserted to be 70% in the aorta and 30% in the ventricle, according to clinical practice. Additionally, the cylindrical stent model is radially translated to account for the patient’s calcification pattern. All the previous steps are detailed in [29]. Moreover, the virtual stent insertion procedure was qualitatively validated in [28] for two patients with available CTA scans taken few years after the TAVI intervention.

Regarding the stent model, an advancement of this work with respect to [29, 28] is the introduction of a wire-frame design in the aortic side of the stent. Specifically, we generate realistic CAD models of the SAPIEN valves’ stent with *SolidWorks* and *Autodesk Fusion* (Figure 1 center). Then, starting from a hollow cylinder, we manipulate its geometry to match the wire-frame design of the CAD model only on the aortic side (Figure 1 right). Notice that the ventricular side of the stent is modeled as a hollow cylinder due to the presence of the inner skirt in the SAPIEN valves (see Figure 1 left).

As for the bio-prosthetic leaflets, two advancements with respect to [29, 28] are the use of a realistic geometry, which mimics the SAPIEN valves’ leaflets [34] (Figure 2), and the modeling of the opening and closure dynamics. An open and a closed configuration of the leaflets are obtained with a mechanical simulation in *Abaqus*. Specifically, the leaflets are initially modeled in a stress-free configuration. A physiological pressure waveform is then applied to the leaflet surfaces, while the leaflet-to-stent attachment region is fully constrained. Following the simulation of an entire cardiac beat, the fully open and closed configurations of the bioprosthetic valve leaflets are exported as STL files for the subsequent step of the workflow. Then, we compute a distance from the closed to the open configuration and we use it to describe the opening and closure dynamics (see Section 2.4). Finally, the leaflets geometry is inserted in the stent model so that it has the same orientation of the stent and it is consistent with its wire-frame design.

**Figure 2:**
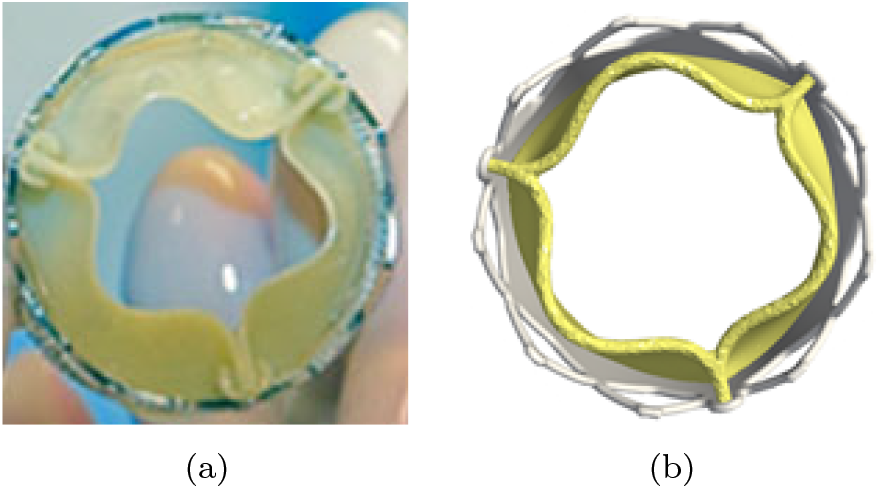
SAPIEN leaflets in open configuration: image of the real valve (a) and model used in the CFD simulations (b). The image of the real valve is taken from Holoshitz, Kavinsky, Hijazi [35] with written permission by Hijazi.

The resulting computational domain, depicted in Figure 3a, has one inlet and one outlet sections. Additionally, the external wall comprises the interface between blood and the stent, the interface between blood and the native annulus and the aortic wall.

**Figure 3:**
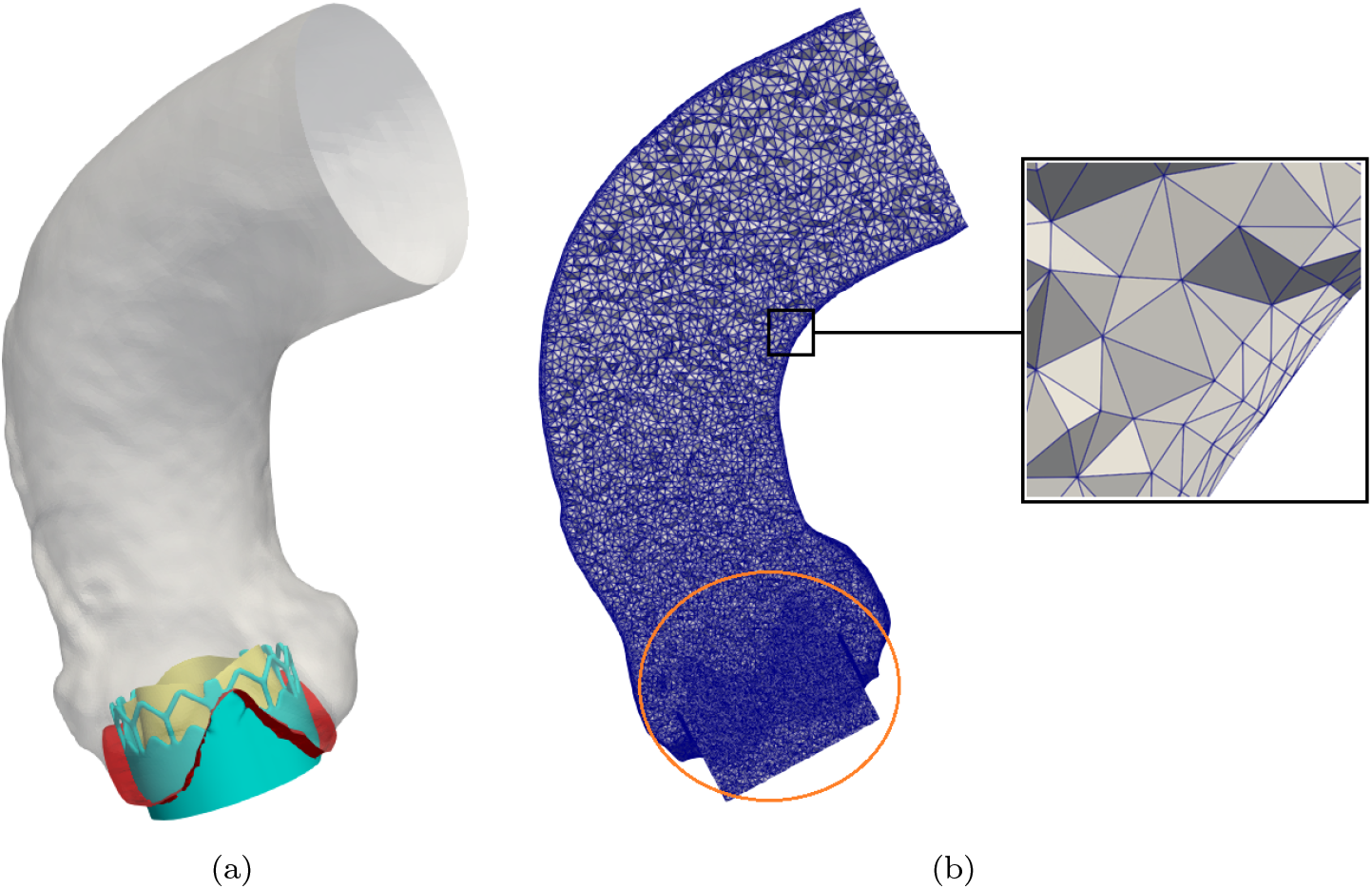
(a) Computational domain comprising the interface between blood and the stent (light blue), the interface between blood and the native annulus (red), the aortic wall (gray), and the leaflets geometry (yellow). (b) Volumetric computational mesh with a boundary layer (zoom in black box) and a local refinement in the leaflet area (highlighted in orange).

As a last step we generate the computational mesh made of tetrahedral elements which are finer close to the inlet and coarser going toward the outlet. On the whole aortic wall we consider a boundary layer made of three sublayers with thicknesses {0.1, 0.2, 0.4} *mm* going from the wall to the internal domain. Moreover, we refine the mesh to a 0.25 *mm* size in the leaflets area to ensure a proper implicit representation of the leaflets in the numerical simulations (see Section 2.4). The resulting mesh has an average size of ≈ 0.8 *mm* and is chosen after a mesh sensitivity analysis (see Appendix A)

### 2.3. Personalized flow rate profile

In accordance with [28], we want to prescribe a flow rate condition at the inlet section of the computational domain (Figure 3a). In particular, in a physiological flow rate profile was taken from literature [36] and its magnitude was scaled to match the patient’s cardiac output. In order to further personalize our CFD model, in this work we assume that the inlet flow rate shares the same temporal evolution profile as the blood velocity inside the Left Ventricular Outlet Tract (LVOT), reconstructed from the available PW Doppler TTE image. The resulting flow rate is characterized by a patient-specific mean value (i.e. cardiac output), heart rate, and partition between systolic and diastolic phase.

More specifically, PW Doppler TTE images show the temporal evolution of the blood velocity at the center of the LVOT during few heartbeats. For each heartbeat, we manually draw the velocity temporal evolution during systole, following the Doppler signal’s contour and the drawings performed by clinicians, if present. These manual drawings were validated by an expert clinician for the whole study population. Then, using *WebPlotDigitizer*, we digitalize the drawn profiles and we identify the end of the heartbeat. In particular, the end of the heartbeat is identified exploiting the electrocardiography signal, available in the image, and the start of the velocity signal of the following heartbeat. Finally, we average the digitalized signals between the heartbeats and we scale the averaged profile to match the patient-specific cardiac output and heart rate. The full procedure is schematized in Figure 4.

**Figure 4:**
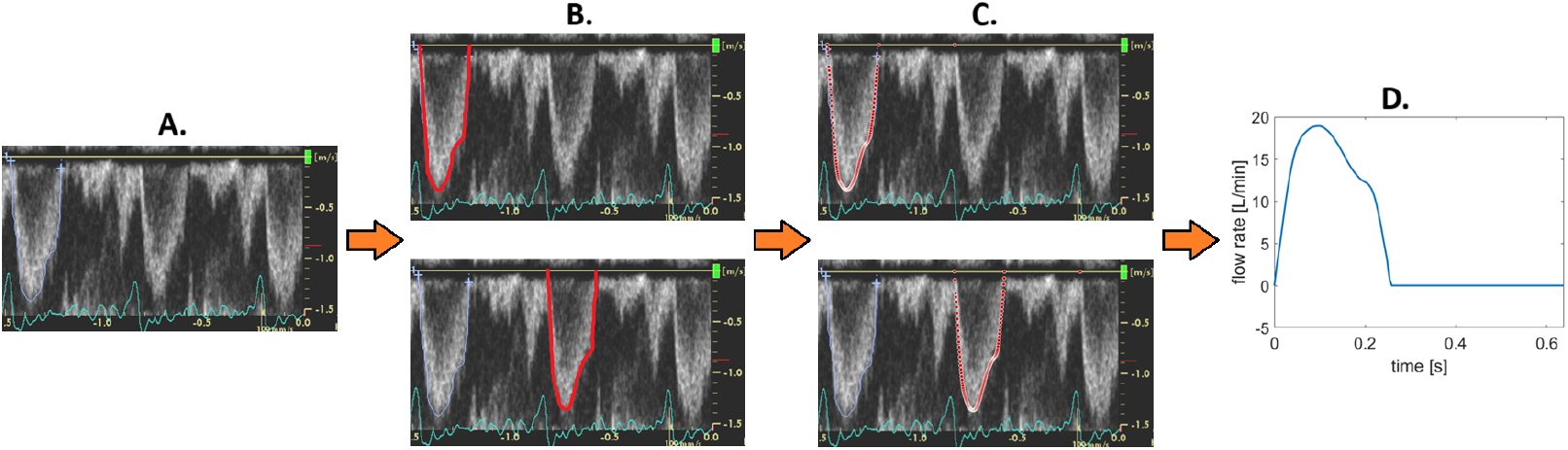
Derivation of the personalized flow rate profile. Starting from a PW Doppler TTE image (A.), we manually draw the temporal evolution of the blood velocity during systole for each visible heartbeat (B.), we digitalize each drawing and identify the end of the heartbeat (C.), we average the digitalized profiles between the heartbeats and we rescale the averaged profile to match the patient’s cardiac output and heart rate (D.).

Let us notice that, in the clinical setting, the cardiac output is usually computed from PW Doppler TTE data assuming a flat velocity profile in the LVOT. This tends to overestimate the real cardiac output by about 15% [37] due to the skewness of the real velocity profile. Accordingly, in this work we rescale the clinical cardiac output so to eliminate this overestimation (for simplicity assumed to be equal to 15% in all the patients).

### 2.4. Mathematical and numerical modeling

In this work we considered the same mathematical model presented in [28], where the system of equations is reported. In particular, we model blood as an incompressible homogeneous Newtonian fluid with the Navier-Stokes (NS) equations, as commonly done in large vessel like the aorta [38, 39]. We account for the presence of turbulence in the ascending aorta [40] by employing a Large Eddy Simulation (LES) method, which is most suitable due to its capability to capture the laminar, transitional and turbulence features characterizing the cardiac cycle [41]. Specifically, we use the *σ*-LES model [42], which proved to perform well in closed channel configurations, showing accurate results in the near wall region [43]. We provide an implicit representation of the bio-prosthetic leaflets with the Resistive Immersed Implicit Surface (RIIS) method [44]. This method adds to the NS momentum equation a local penalization term which surrogates the obstruction to the flow given by the presence of the leaflets. As for the boundary conditions, we consider the external wall as rigid (CFD approach), we impose a physiological pressure at the outlet section and a flow rate condition, prescribing the personalized flow rate profile presented in Section 2.3, at the inlet section. The latter condition does not provide enough information to close the NS problem, thus it needs to be completed. In order not to set a priori a spatial profile for the velocity field at the inlet, which would affect the accuracy of the numerical solution [45], we prescribe this condition using a Lagrange multiplier [46]. This approach assumes the traction to be normally directed and constant at the inlet section and results in an augmented formulation of the NS systems.

Regarding the modeling of the leaflets opening and closure dynamics, in the numerical simulations we use the distance function introduced in Section 2.2 to interpolate between the closed and open configurations of the leaflets, according to the following law: the opening and closure times are 15 *ms* and 45 *ms*, respectively, according to [47]; moreover, the transition between the two configurations is regulated by a valve opening coefficient which follows the cosine-exponential ramps

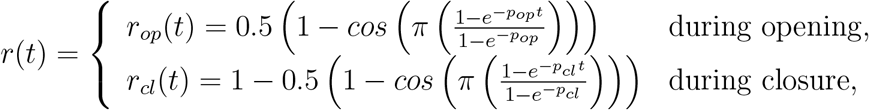

where *t* is a standardized time and *p* = {*p*_*op*_, *p*_*cl*_ } is a parameter to be calibrated for each phase. We vary *p*_*op*_ ∈ [2.5, 3.5] for the opening ramp and *p*_*cl*_ ∈ [ −3, −2] for the closure ramp according to the slope of the input flow rate profile at the opening and closure instants. Specifically, patients showing steeper flow rate profiles are associated to higher absolute values of *p*. We stress that the leaflets geometry is a surface made of shell elements, we implicitly represent it as an immersed surface and provide a thickness to it directly in the numerical simulations using the RIIS method (see [28] for details).

Let us now present the numerical methods employed in this study. A spatial discretization of the problem is achieved using a Finite Element approach with piece-wise linear elements both for the velocity and pressure fields, referred to as P1-P1. A Streamline Upwind Petrov–Galerkin/Pressure Stabilizing Petrov–Galerkin (SUPG/PSPG) stabilization [48] is considered to account for the advection-dominated regime and the use of P1-P1 spaces, which are inf-sup unstable. Possible numerical instabilities arising from the presence of recirculating areas at the outlet section are addressed with a backflow stabilization [49]. Moreover, the problem is discretized in time with a first order backward Euler scheme with a semi-implicit treatment of the convective term and an explicit treatment of the sub-grid scale viscosity, introduced by the *σ*-LES method. We consider a different time-step for each patient according to the duration of the patient’s systolic phase. Specifically, a time-step Δ*t* = 10^−3^ *s* is used for patient NODEG6 who shows the longest systolic period (0.4 *s*) and for the other patients the time-step is scaled so that we have roughly the same number of time instants during systole. Finally, the augmented algebraic system arising from the introduction of a Lagrange multiplier to impose the inlet flow rate condition is monolithically solved using a preconditioned GMRES iterative solver [50], exploiting an ad-hoc preconditioner [51].

For the implementation of the mathematical and numerical methods presented in this subsection we rely on the multi-physics high-performance Finite Element library life^x^ [52, 53], developed at MOX, Dipartimento di Matematica, with the collaboration of LaBS, Dipartimento di Chimica, Materiali ed Ingengneria Chimica (both at Politecnico di Milano).

### 2.5. Post-processing of the results

Due to the turbulent nature of the analyzed flow, in this paper we numerically simulate eight complete heartbeats, discarding the first two to avoid the influence of the null initial condition. In particular, using the *Paraview* software, we post-process the *phase-average* (also named as *ensemble*) velocity and pressure fields, i.e., the average at each time over the six heartbeats:

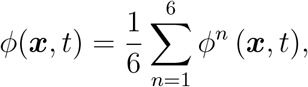

where *ϕ*^*n*^ is either the velocity or the pressure at the *n*-th heartbeat. Specifically, in this paper we aim to propose computational hemodynamic indices, obtained by the phase-average fields, in early post-TAVI scenarios that correlate with a premature onset of SVD, detected at 5-10 years follow-up exam.

#### 2.5.1. Quantities of interest

Let us define the fluid traction on a surface as:

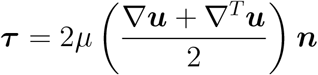

where *µ* is the blood viscosity, ***u*** is the (phase-average) blood velocity and ***n*** is the surface normal. Accordingly, the Wall Shear Stress (WSS) vector reads:

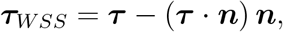

which allows us to define the following quantities:

- Time-Averaged WSS (*TAWSS*):

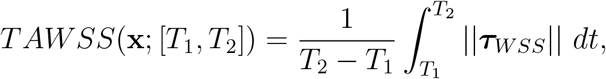

where *T*_1_ and *T*_2_ identify the time-period (systole, diastole or complete heartbeat) which could change according to the index, and ||***v***|| is the magnitude of a generic vector ***v***;
- Oscillatory Shear Index (*OSI*):

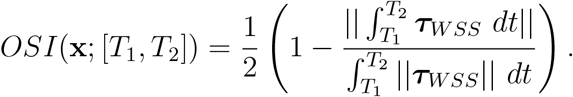

It provides a measure of the oscillatory nature of the WSS vector;
- Topological Shear Variation Index (*TSV I*):

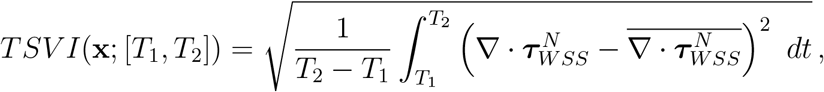

where ∇· is the divergence operator, 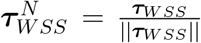 the normalized WSS vector, and 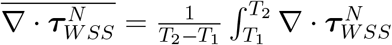 *dt* the average in time of the divergence of the normalized WSS vector. It represents the variation in the WSS action on the vessel expansion/contraction [54].

#### 2.5.2. Hemodynamic indices and scores

SVD has been associated with an increased infiltration of host cells (e.g. immune cells, cell debris) in the bio-prosthetic leaflets [8, 9, 26]. Accordingly, we propose three computational hemodynamic indices in order to study their correlation with a premature onset of SVD.

A diffuse damage/inflammation of the aortic wall can lead to the recruitment of immune cells to the aortic root [55], thus facilitating their infiltration in the leaflets. To quantify this damage we propose the Wall Damage Index (*WDI*) computed on the aortic wall:

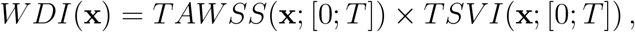

where *T* is the duration of the whole heartbeat. It is known that high WSS, here associated with high *TAWSS* values, on the vessel wall can result in endothelial changes, possibly promoting inflammation [56]. Additionally, significant variations in WSS directionality, corresponding to high *TSV I* values, can affect valvular diseases by influencing endothelial cells [57]. Thus, we associate high *WDI* values with a possible damage of the aortic wall.

High stresses on the bio-prosthetic leaflets can cause the delamination of the connective tissue, promoting the infiltration of immune cells [7]. For this reason, we propose the Leaflet Delamination Index (*LDI*) computed on the ventricular side of the leaflets:

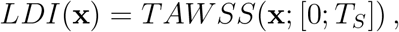

where *T*_*S*_ is the systolic end instant.

Low and oscillatory WSS is known to impair endothelial integrity, thus increasing its permeability [58]. Accordingly, we propose the Leaflet Permeability Index (*LPI*) computed on the aortic side of the leaflets:

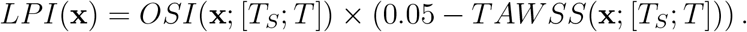

Notice that, high *OSI* and low *TAWSS* values result in high *LPI* values.

To investigate the correlation between the proposed indices and a premature onset of SVD, we define a hemodynamic score associated to each index by spatially averaging the index on the corresponding surface. Specifically, *WDI* is averaged on the whole aortic wall to obtain 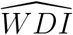, *LDI* is averaged on the ventricular side of the leaflets to obtain 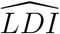, and *LPI* is averaged on the aortic side of the leaflets to obtain 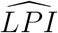.

#### 2.5.3. Synthetic score

With the aim of clearly separating between patients with and without SVD at long-term follow-up exam (DEG vs NODEG), we define a global synthetic score, which combines the proposed hemodynamic scores.

In order to be combined, the hemodynamic scores have to first be standardized. To do so, similarly to [28], we compute the 25*th* percentile (*Q*_1_), the 75*th* percentile (*Q*_3_), and the inter-quartile range (*IQR* = *Q*_3_ − *Q*_1_) for each hemodynamic score in the whole study population and we linearly project the interval [*Q*_1_ − *IQR, Q*_3_ + *IQR*] into the interval [−1, 1]. The standardization procedure is performed using the *Matlab* software.

The global synthetic score *SV D*_*score*_ reads:

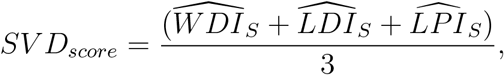

where the subscript *S* identifies the hemodynamic standardized score.

#### 2.5.4. Validation of the numerical results

To validate the numerical results of our CFD model we use the maximum blood velocity measurements through the bio-prosthetic valve taken from early post-TAVI CW Doppler TTE data, available for all patients in the study population (apart from DEG2). During CW Doppler TTE a transducer continuously transmits ultrasound waves and receives echoes in order to measure blood velocities along an entire beam Γ oriented as the valve and passing through its center, see Table 1-right. The maximum blood velocity through the valve is identified as the maximum velocity magnitude obtained along the beam Γ during the heartbeat, and we compare it with the same quantity extracted from the CFD results, assuming a Doppler-like beam Γ with a radius of 1 *mm*.

**Table 1:** Left: Numerical blood velocity validation. Early post-TAVI maximum blood velocity magnitude through the bio-prosthetic valve taken from CW Doppler TTE data 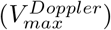 and from the CFD results 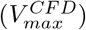, and relative error between the two measures (*Err*). The error range is computed accounting for the decimal truncation of 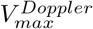 (i.e. ±0.05 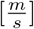). Right: Visualization of the Doppler image and an example of the region Γ where the maximum blood velocity magnitude is computed from the CFD results.

| ID | $V_{max}^{Doppler} [\frac{m}{s}]$ | $V_{max}^{CFD} [\frac{m}{s}]$ | $Err [\%]$ |
| --- | --- | --- | --- |
| DEG1 | 2.6 | 2.92 | 10.6-14.5 |
| DEG4 | 2.6 | 2.51 | 1.6-4.9 |
| DEG5 | 2.5 | 2.33 | 4.9-8.2 |
| DEG6 | 2.6 | 2.62 | 0.0-2.7 |
| DEG7 | 2.4 | 2.39 | 0.0-2.0 |
| NODEG1 | 2.5 | 2.54 | 0.0-3.7 |
| NODEG2 | 2.0 | 2.04 | 0.0-4.6 |
| NODEG4 | 2.5 | 2.51 | 0.0-2.4 |
| NODEG5 | 2.3 | 1.85 | 17.8-20.9 |
| NODEG6 | 2.1 | 2.29 | 7.0-11.7 |
| NODEG7 | 2.5 | 2.08 | 15.1-18.1 |

## 3. Results

All the numerical simulations were run in parallel on 112 cores of Intel Sapphire Rapids@2.00GHz CPU’s, using the computational resources provided by the CINECA supercomputing center, Italy. The average computational time for each simulation (i.e. for eight complete heartbeats) is 17.0 ± 5.3 hrs.

### 3.1. Outcomes of the validation of the numerical results

In table 1 we report the maximum blood velocity obtained from CW Doppler TTE and from the CFD results (see Section 2.5.4). Out of the eleven patients with available clinical datum, the table shows that: in four patients (DEG6, DEG7, NODEG1, NODEG4) the agreement is excellent and the error is very close to zero; in two patients (DEG4, NODEG2) there is very slight increase of the error, which however is less than 5%; in other two patients (DEG5, NODEG6) we have a good agreement with errors around 10%; in the remaining three patients (DEG1, NODEG5, NODEG7) we experienced increased errors up to 20%.

### 3.2. Analysis of the hemodynamic patterns

In the following we analyze the CFD results for patient DEG1, as a representative of the study population.

In Figure 5 we report, for three selected instants during systole^1^, the CFD velocity streamlines and pressure on a longitudinal plane, and the WSS magnitude patterns on the aortic wall. From the velocity plots we can observe that the presence of a small size TAVI valve (23 *mm* in external diameter) gives rise to the a high-velocity jet forming across the implanted valve, together with some vortical structures along side the jet (early systole). Let us note that, the velocity magnitude obtained inside the jet strictly depends on the personalized flow rate profile (see Section 2.3) prescribed at the inlet, since the geometric opening area of the valve is the same for each patient in our model. The impingement of the vortices on the aortic wall (peak systole) results in very chaotic hemodynamic patterns with recirculating areas in the ascending aorta (late systole). Additionally, the accelerated flow results in significant pressure gradients across the valve, with maximum values around 20 [*mmHg*]. In particular, in the first two time instants, low pressure values are obtained near the vortices inside the ascending aorta, with a pressure recovery towards the outlet section. Furthermore, high WSS magnitudes are obtained on the aortic wall due to the collision of the jet on the wall.

**Figure 5:**
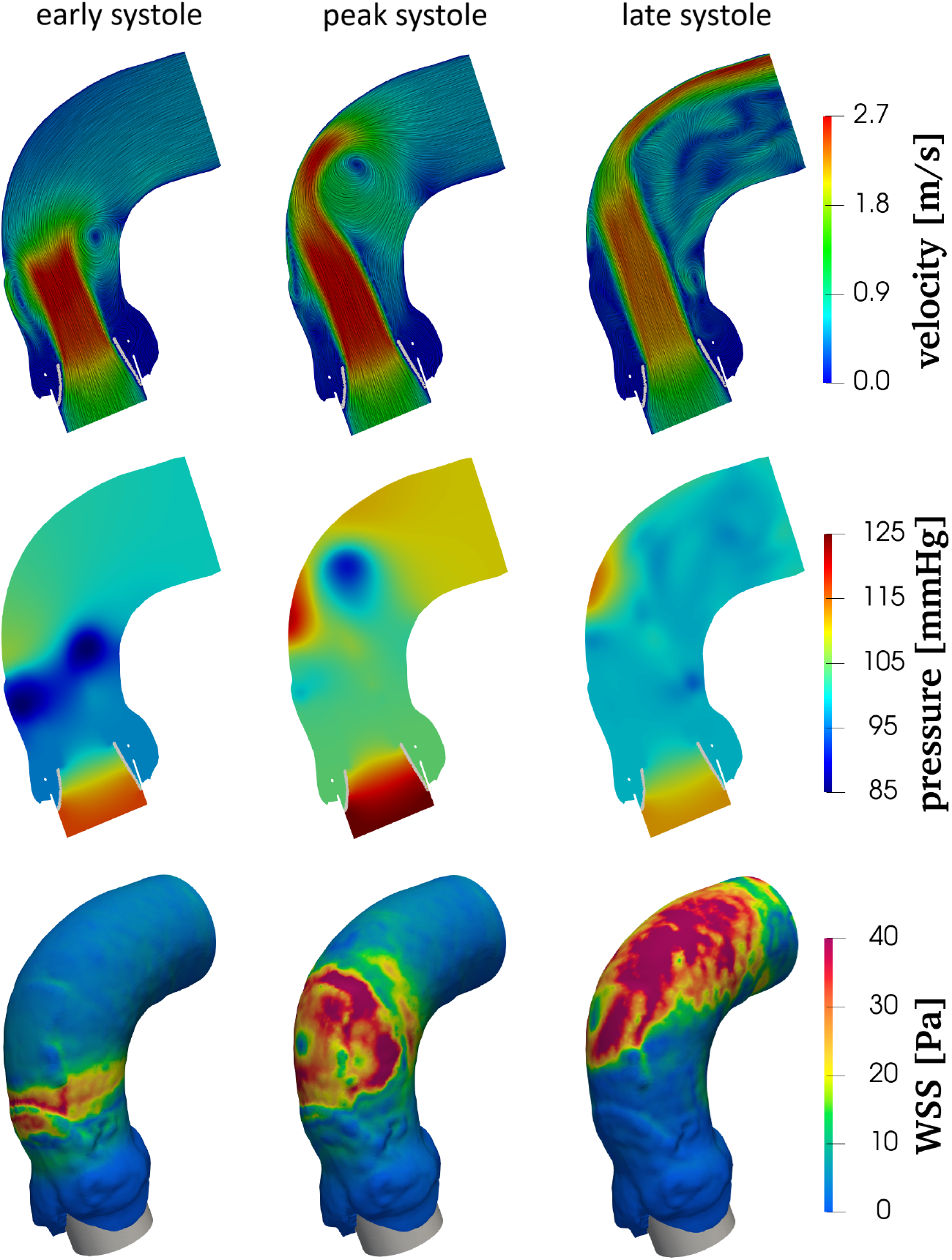
Evolution of the velocity streamlines (top) and pressure (middle) on a longitudinal plane, and of the WSS magnitude on the the aortic wall (bottom) at three selected instants during systole (for the represented patient DEG1, they are *t* = 0.063 *s, t* = 0.093 *s, t* = 0.159 *s*).

In Figure 6 we report the spatial distribution of the hemodynamic indices presented in Section 2.5.2, which is driven by the complex hemodynamics described above. Indeed, the high stresses on the aortic wall and the disturbed flow patterns, leading to significant topological variations of the WSS vectors, are the main determinants for the *WDI* distribution. Furthermore, the accelerated flow across valve affects the stress intensity on the ventricular side of the leaflets during systole, thus the *LDI*. Finally, the chaotic flow patterns, characterized by several vortical structures in the aorta, leads to low and oscillatory stresses on the aortic side of the leaflets during diastole, thus influencing the *LPI* intensity and patterns.

**Figure 6:**
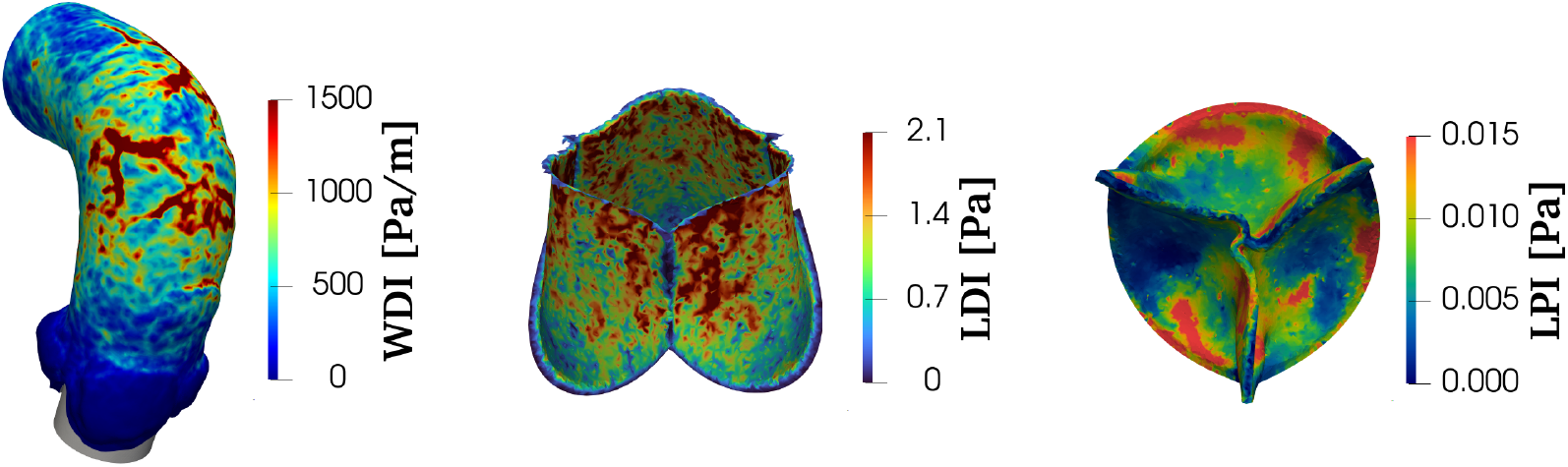
Hemodynamic indices *WDI, LDI* and *LPI* (see Section 2.5.2). Patient DEG1.

### 3.3. Hemodynamic and synthetic scores

In Table 2 we report the values of each hemodynamic score (see Section 2.5.2) in the study population. From the table it is clear that different scores take value in significantly different ranges, thus stressing the importance of the standardization procedure, described in Section 2.5.3, in order to combine them.

**Table 2:** Hemodynamic scores 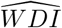, 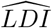 and 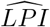 for each patient.

| ID | $\widehat{WDI} [\frac{Pa}{m}]$ | $\widehat{LDI} [Pa]$ | $\widehat{LPI} [Pa]$ |
| --- | --- | --- | --- |
| DEG1 | $5.10 \times 10^2$ | $1.08 \times 10^0$ | $7.27 \times 10^{-3}$ |
| DEG2 | $5.31 \times 10^2$ | $1.12 \times 10^0$ | $5.58 \times 10^{-3}$ |
| DEG4 | $4.03 \times 10^2$ | $1.12 \times 10^0$ | $7.49 \times 10^{-3}$ |
| DEG5 | $5.28 \times 10^2$ | $1.07 \times 10^0$ | $4.81 \times 10^{-3}$ |
| DEG6 | $5.83 \times 10^2$ | $1.24 \times 10^0$ | $6.11 \times 10^{-3}$ |
| DEG7 | $4.50 \times 10^2$ | $1.17 \times 10^0$ | $6.14 \times 10^{-3}$ |
| NODEG1 | $3.98 \times 10^2$ | $1.06 \times 10^0$ | $3.36 \times 10^{-3}$ |
| NODEG2 | $2.69 \times 10^2$ | $0.81 \times 10^0$ | $4.64 \times 10^{-3}$ |
| NODEG4 | $3.85 \times 10^2$ | $0.98 \times 10^0$ | $7.14 \times 10^{-3}$ |
| NODEG5 | $3.07 \times 10^2$ | $0.82 \times 10^0$ | $5.29 \times 10^{-3}$ |
| NODEG6 | $4.31 \times 10^2$ | $0.99 \times 10^0$ | $4.39 \times 10^{-3}$ |
| NODEG7 | $5.03 \times 10^2$ | $0.95 \times 10^0$ | $3.11 \times 10^{-3}$ |

In Figure 7 we plot the hemodynamic scores distributions in the DEG and NODEG sub-populations using boxplots. The figure shows that each score tends to be characterized by greater values for DEG patients with respect to NODEG patients. In order to properly assess the observed trend, for each individual hemodynamic score we employ, using *Matlab*, a right-tailed Wilcoxon rank-sum test having as alternative hypothesis that the median of the DEG distribution is statistically greater than the median of the NODEG distribution. This non-parametric test is chosen due to the limited amount of observations. The p-values resulting from these tests are: *p* = 0.0076 for 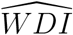, *p* = 0.0011 for 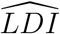, and *p* = 0.0206 for 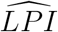. Thus, with respect to each hemodynamic score the DEG distribution is statistically greater than the NODEG distribution at a 2.5% significance level.

**Figure 7:**
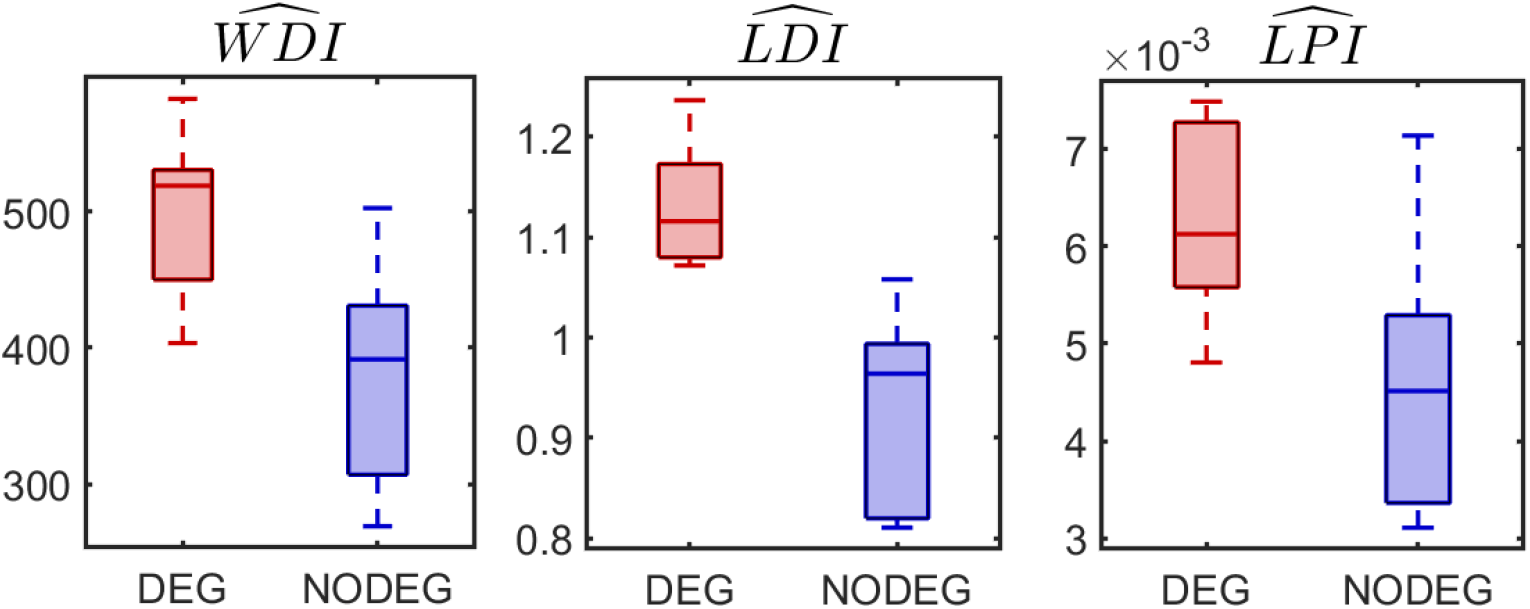
Boxplot distributions of each hemodynamic score in the DEG and NODEG sub-populations.

Lastly, in Table 3 we report the global synthetic score *SV D*_*score*_ (see Section 2.5.3) in the whole population. The *SV D*_*score*_ is able to clearly separate between the two sub-populations, with each DEG patient showing positive values and each NODEG patient showing negative values of this score.

**Table 3:** Global synthetic score *SV D*_*score*_ for each patient.

| ID | $SVD_{score}$ | ID | $SVD_{score}$ |
| --- | --- | --- | --- |
| DEG1 | 0.33 | NODEG1 | −0.30 |
| DEG2 | 0.24 | NODEG2 | −0.76 |
| DEG4 | 0.22 | NODEG4 | −0.04 |
| DEG5 | 0.09 | NODEG5 | −0.61 |
| DEG6 | 0.57 | NODEG6 | −0.23 |
| DEG7 | 0.24 | NODEG7 | −0.31 |

## 4. Discussion

The mechanisms underlying the development of SVD in transcatheter bio-prosthetic valves are still not fully understood: multiple active processes are involved, from mechanical stresses to long-term immune rejection and atherosclerosis-like tissue remodeling [7]. Specifically, inflammatory reaction and immune response may play a crucial role in the pathogenesis of SVD, which is associated with an elevated immune cell infiltration in the bio-prosthetic leaflets [8, 9]. Clinical studies identified some risk factors related to SVD, such as small prosthesis size [11], a valve-in-valve procedure [10], young age, patient-prosthesis mismatch, larger body surface area, and smoking [12]. Also, in [13] the authors hypothesized that an accelerated flow across the implanted valve, due to a high cardiac output, could be a factor in the development of SVD.

In this context, computational modeling may provide a detailed description of post-TAVI hemodynamics allowing to investigate the accelerated flow features possibly influencing SVD.

Accordingly, in this study we introduced new hemodynamic computational risk factors for the premature onset of SVD. Specifically, we proposed three hemodynamic scores (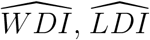, and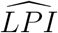), that can be related to an infiltration of host cells in the bio-prosthetic valve leaflets (see Section 2.5.2), and a global synthetic score (*SV D*_*score*_) which combines such scores (see Section 2.5.3). This is done by numerically simulating post-TAVI hemodynamics in a cohort of patients for which a follow-up at 5-10 years is available.

First, we notice that in our study we analyzed post-TAVI hemodynamics just after the implantation. This is based on the assumption that the hemodynamic patterns initiating the development of SVD are influential already a short time after the intervention [31], when the implanted bio-prosthetic valve is still fully functioning. This allowed us to avoid the modeling of the bio-prosthetic leaflets degeneration process.

In order to try to isolate the effect of hemodynamics on the development of SVD, we considered TAVI valves with very similar designs (Edwards SAPIEN of first and second generation). Indeed, different valve designs can influence valve deterioration due, for example, to the different distribution of mechanical stresses [25] and the different material properties of the leaflets [59]. Future studies could focus on different valve designs to investigate if the trends identified here still hold true.

Regarding the personalized flow rate profile (Section 2.3) prescribed in the numerical simulations at the inlet, we assumed that it shares the same temporal trend of the blood velocity in the LVOT, measured for the patient by Echo Doppler. The accuracy of this assumption depends on the skewness of the velocity profile, which is not known. However, the phase discrepancy between the two signals is in any case small, and this strategy is anyway patient-specific, thus more accurate than considering a flow rate taken from the literature, as done for example in [28].

In the literature, combinations of *TAWSS* and *OSI* to identify vessels ≈ regions subject to low and oscillatory shear stresses are commonly based on the use of the Endothelial Cell Activation Potential (*ECAP*) [60, 61] and Relative Residence Time (*RRT*) [62], which depend on the inverse of *TAWSS*. Our numerical results are often characterized by values of *TAWSS* ≈ 0 in some very localized areas of the leaflets during diastole, leading to very high *ECAP* and *RRT* values that would dominate over the spatial average needed to the define the hemodynamic score. For this reason, in this study we proposed *LPI* which does not have a singularity for *TAWSS* = 0 (Section 2.5.2).

Doppler-based blood velocity measurements showed a variability up to 9% between different measurements performed by the same experienced vascular technologist in an idealized in-vitro setting [63]. Accordingly, we found a good quantitative agreement between the CW Doppler TTE and CFD measures in the validation analysis reported in Section 3.1, with seven out eleven patients showing errors smaller than the 9% threshold, and patient NODEG6 showing errors around 10% (close to the threshold). The greater discrepancies observed for patients DEG1, NODEG5 and NODEG7 can be mainly due to two reasons: an under/over expansion of the real implanted TAVI valve, that we cannot model since we do not have at disposal any intra-procedural data; an inaccurate computation of the personalized flow rate profile, due to a mismatch between the flow rate and blood velocity temporal trends for these patients or a sub-optimal reconstruction of the velocity temporal profile from the PW Doppler TTE image.

The analysis of the computational hemodynamic patterns (Section 3.2) highlighted the mildly stenotic scenario that follows the implantation of a small size TAVI valve, characterized by high velocity magnitudes and a pressure recovery distal to the aortic valve [64]. This is consistent with the observation that all stented bio-prosthetic valves result in a residual stenosis of up to 20% [65]. Specifically, the formation of a high-velocity jet in the ascending aorta leads to complex and disturbed hemodynamic patterns possibly speeding up SVD pathogenesis through inflammatory reaction and infiltration of host cells in the leaflets.

We identified three hemodynamic indices in early post-TAVI scenarios that showed statistically significant differences between patients with and without SVD at long-term follow-up exam (see Section 3.3). This is a very promising result showing that patients with a prematurely degenerated valve belong to a different statistical distribution with respect to the considered hemodynamic features even when the valve is not degenerated yet. Therefore, this analysis represents a preliminary proof of the influence of aortic hemodynamics on the development of SVD, which should be considered for a proper assessment of SVD and has been rarely investigated with computational models.

The definition of the global synthetic score *SV D*_*score*_ allowed us to obtain a scoring system able to clearly separate patients with and without SVD in view of a possible clinical translation (see Table 3). We found that the global score is more robust than the three individual hemodynamic scores, probably since it takes into account multiple mechanisms facilitating immune cells infiltration (aortic wall damage/inflammation, leaflet delamination, leaflet increased permeability). Additionally, we notice that also the individual 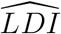 score alone is able to separate the two sub-populations (see Figure 7-middle), even if with a less evident separation (0.13 for *SV D*_*score*_, see Table 3, vs 0.06 for the standardized 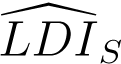, not reported here for the sake of exposition).

The proposed synthetic *SV D*_*score*_ could be easily translated into clinical practice to identify, early after the intervention, those patients at higher risk of a premature onset of SVD. This could drive a patient-specific planning of follow-up exams to closely monitor the high SVD risk patients, trying to avoid their dispersion after the procedure.

Lastly, we notice that the clinical data needed to build the proposed scores (pre-TAVI CTA scans and early post-TAVI Doppler TTE measures) are routinely acquired within the TAVI context. Since the model does not require any ad-hoc or invasive clinical measurement, our framework can be used to perform both prospective and retrospective studies, providing a detailed and personalized description of the main flow features while sparing patients from additional measurements and exams.

## 5. Limitations

We start by noticing that our mathematical and numerical model (Section 2.4) is characterized by some simplifications:

- We modeled blood as a homogeneous Newtonian fluid and the wall as rigid. This can have an influence on hemodynamic features such as the Wall Shear Stress (WSS) [66]. However, we notice that we study elderly patients which are usually characterized by a limited tissue elasticity due to stiffening and calcifications. Moreover, in large vessel like the aorta, the influence of blood cells can be neglected and the Newtonian assumption is the standard one [67];
- We did not account for the coronary branches and thus for the corresponding blood flow. This could affect blood-dynamics inside the aortic root, specially during diastole, and thus the hemodynamic indices proposed in Section 2.5.2, specially *LPI*, being computed during the diastolic phase. Notice that, considering coronary flow with a rigid aortic wall could lead to non-physiological flow patterns since the blood flowing to the coronaries is thought to come from the relaxation of the aortic root wall following its systolic expansion [68]. In any case, we believe that the inclusion of the coronary flow should have a common effect on the hemodynamic quantities among patients;
- We provided an implicit dynamic representation of the bio-prosthetic leaflets with the RIIS method. This allowed us to capture the main flow features, such as the opening/closure dynamics and a realistic opening area. However, a Fluid-Structure Interaction (FSI) approach should be able to represent complex features, such as leaflets fluttering, and results in a more accurate computation of the stresses on the leaflets, thus could be considered in future studies.

These simplifications significantly reduced the computational cost of the numerical simulations to make our model suitable to analyze a great number of patients in a timely manner, in view of a potential clinical translation. Additionally, in this study we are interested in comparing numerical results in different patients rather than obtaining optimally accurate results for each patient. Since a fair comparison between patients is guaranteed by the adoption of the same assumptions, then we accept these simplifications.

Moreover, this study is subject to some further main limitations:

- We virtually inserted the bio-prosthetic valve in the aortic geometries assuming a perfect implantation, with the stent obtaining its nominal circular cross-section and being perfectly sealed to the aortic annulus (Section 2.2). An improvement could be achieved by numerically simulating the implantation of the valve using a structural Finite Element Analysis. This allows to obtain the deformation of the stent shape due to the deployment in a calcified valve and to capture paravalvular leakage. However, we accept this simplification since balloon-expandable valves are expected to achieve an almost cylindrical shape after implantation [69], as shown from the post-TAVI CTA scans reconstructions in [28], and since we are mainly interested in studying SVD rather than other complications related to the TAVI procedure;
- The significance of our findings is limited by the sample size of our study population. Therefore, enlarging the study population is crucial to move toward clinical relevance.

## Glossary

TAVI: Transcatheter Aortic Valve Implantation
SVD: Structural Valve Deterioration
CFD: Computational Fluid Dynamics
CTA: Computed Tomography Angiography
PW: Pulsed Wave
TTE: Transthoracic Echocardiography
CW: Continuous Wave
LVOT: Left Ventricular Outlet Tract
NS: Navier-Stokes
LES: Large Eddy Simulation
RIIS: Resistive Immersed Implicit Surface
WSS: Wall Shear Stress
TAWSS: Time-Averaged Wall Shear Stress
OSI: Oscillatory Shear Index
TSVI: Topological Shear Variation Index
WDI: Wall Damage Index
LDI: Leaflet Delamination Index
LPI: Leaflet Permeability Index
ECAP: Endothelial Cell Activation Potential
RRT: Relative Residence Time

## Appendix A. Mesh sensitivity analysis

In this appendix we present the sensitivity analysis performed to choose the mesh size of our computational mesh (Figure 3b). This is done to ensure that our numerical results do not strongly depend on the chosen mesh while allowing for a sustainable computational cost of the simulations. The analysis is carried out only for patient DEG1, as a representative of the study population, and we numerically simulate two complete heartbeats, analyzing only the results of the second one.

In our computational mesh the elements size is minimum at the level of the aortic annulus and a maximum at the level of the outlet section. We select the minimum element size on the aortic wall as a characteristic mesh size *H* and perform sequential refinements of the mesh by decreasing the value of *H*, while keeping the same ratio between the maximum and minimum element sizes. Specifically, we consider three refinement steps associated to {*H*_1_, *H*_2_, *H*_3_ } = {0.82, 0.65, 0.52} *mm*, corresponding to computational meshes {*M*_1_, *M*_2_, *M*_3_ }, respectively. Let us define the refinement factor between two consecutive meshes as 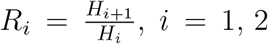. We deem a mesh as sufficiently refined if the relative difference between quantity of interest *Q*_*i*+1_, computed in *M*_*i*+1_, and *Q*_*i*_, computed in *M*_*i*_, is less than a tolerance *T* (*Q, R*_*i*_) which depends on the quantity of interest and the refinement factor. In particular, analogously to [70], we define the tolerance as follows:

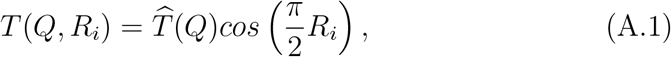

where 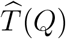 represents an acceptable error with respect to the real measured quantity *Q* in the limit *R*_*i*_ → 0, e.g. the uncertainty related to a clinical measure. This framework stresses the importance of taking into account the amount of refinement between meshes when quantifying the tolerance, with values of *R*_*i*_ ≈ 1 resulting in *T* (*Q, R*) ≈ 0.

We are interested in studying the WSS on the aortic wall, which highly depends on the mesh size. Thus, let us take as a quantity of interest the average-in-time maximum WSS value on the aortic wall 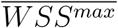 and set 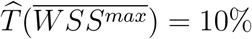, since accurately computing the WSS is challenging even with the most sophisticated imaging techniques. We report in Table A.4 the results of our analysis, showing that mesh *M*_2_ is appropriately refined for our study. For the sake of completeness, in Figure A.8 we plot the WSS on the aortic wall at the peak systolic flow instant for all the meshes, highlighting the similar results obtained for meshes *M*_2_ (*H* = 0.65 *mm*) and *M*_3_ (*H* = 0.52 *mm*).

**Figure A.8:**
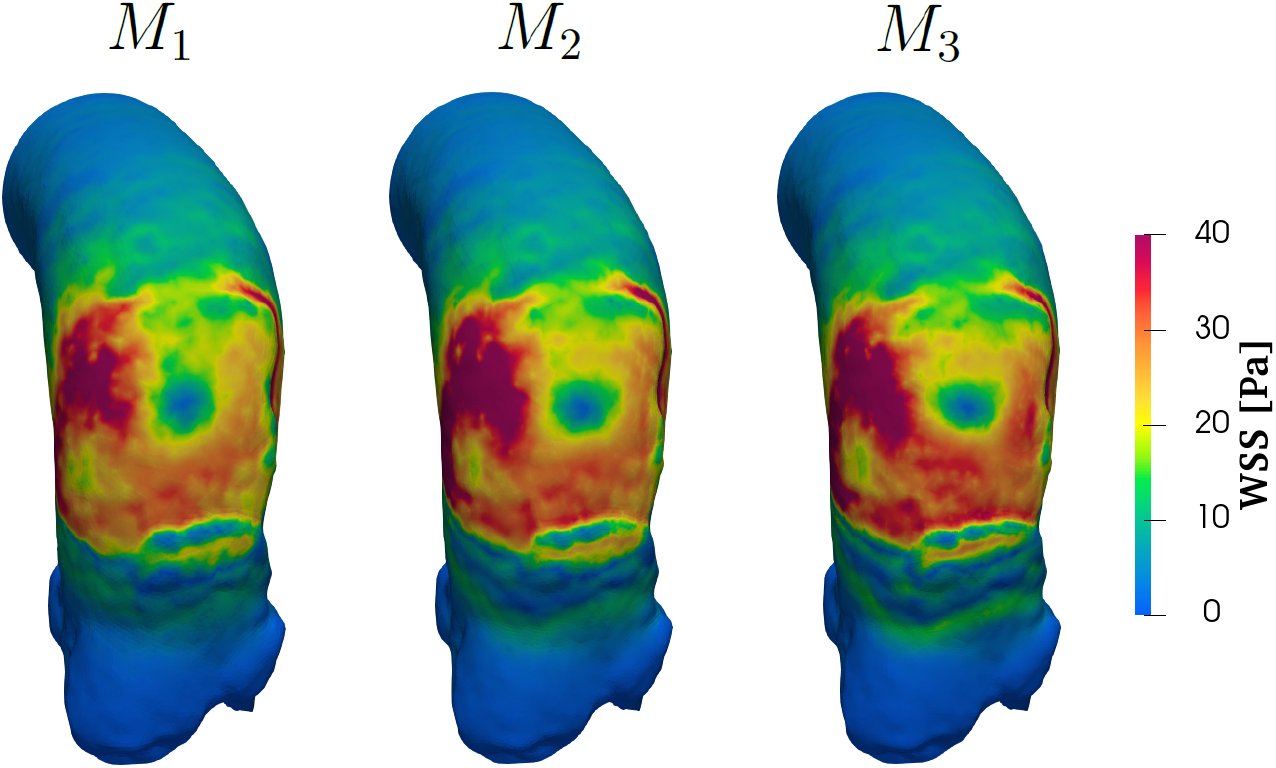
WSS on the aortic wall at the peak systolic flow instant in meshes *M*_1_ (*H*_1_ = 0.82 *mm*, left), *M*_2_ (*H*_2_ = 0.65 *mm*, center) and *M*_3_ (*H*_3_ = 0.52 *mm*, rigth)

**Table A.4:** Mesh sensitivity analysis. i: refinement step; *H*_*i*_: minimum element size on the aortic wall; *R* : refinement factor; 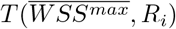: tolerance; 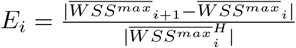 relative difference.

| $i$ | Mesh | $H_i$ | $R_i$ | $T(\overline{WSS}^{max}, R_i)$ | $E_i$ |
| --- | --- | --- | --- | --- | --- |
| 1 | $M_1$ | $0.82\text{ mm}$ | 0.79 | 3.24% | 10.64% |
| 2 | $M_2$ | $0.65\text{ mm}$ | 0.80 | 3.09% | 1.16% |
| 3 | $M_3$ | $0.52\text{ mm}$ | | | |

## Acknowledgments and declarations

Luca Crugnola and Christian Vergara are members of the INdAM group GNCS “Gruppo Nazionale per il Calcolo Scientifico” (National Group for Scientific Computing). CV has been partially supported by: i) the European Union-Next Generation EU, Mission 4, Component 1, CUP: D53D23018770001, under the research project MIUR PRIN22-PNRR n.P20223KSS2, “Machine learning for fluid structure interaction in cardiovascular problems: efficient solutions, model reduction, inverse problems”, ii) the Italian Ministry of Health within the PNC PROGETTO HUB LIFE SCIENCE - DIAGNOSTICA AVANZATA (HLS-DA) “INNOVA”, PNCE3-2022-23683266–CUP: D43C22004930001, within the “Piano Nazionale Complementare Ecosistema Innovativo della Salute” - Codice univoco investimento: PNCE3-2022-23683266; iii) the Italian research project MIUR PRIN22 n.2022L3JC5T “Predicting the outcome of endovascular repair for thoracic aortic aneurysms: analysis of fluid dynamic modeling in different anatomical settings and clinical validation”; iv) Italian Ministry of Health within the project “CAL.HUB.RIA” - CALABRIA HUB PER RICERCA INNOVATIVA ED AVANZATA. Code: T4-AN-09, CUP: F63C22000530001.

We acknowledge the CINECA award under the ISCRA B and ISCRA C initiatives, for the availability of high-performance computing resources and support (ISCRA grants: IscrB_HPCHeart, P.I. Alfio Quarteroni; IscrC_TAVICFD, P.I. Luca Crugnola; IscrC_compTAVI, P.I. Luca Crugnola).

## Ethics approval

The study was approved from the Institutional Review Board of Centro Cardiologico Monzino and registered with number R1264/20. Informed consent was obtained from all patients.

## Footnotes

1 Notice that such values could be slightly different among patients according to the duration of the corresponding heartbeat.

## References

[1] A. Vahanian, F. Beyersdorf, F. Praz, M. Milojevic, S. Baldus, J. Bauersachs, D. Capodanno, L. Conradi, M. De Bonis, R. De Paulis, V. Delgado, N. Freemantle, M. Gilard, K. H. Haugaa, A. Jeppsson, P. Jüni, L. Pierard, B. D. Prendergast, J. R. Sádaba, C. Tribouilloy, W. Wojakowski, E. S. D. Group, E. N. C. Societies, 2021 ESC/EACTS Guidelines for the management of valvular heart disease: Developed by the Task Force for the management of valvular heart disease of the European Society of Cardiology (ESC) and the European Association for Cardio-Thoracic Surgery (EACTS), European Heart Journal 43 (2021) 561–632.

[2] C. M. Otto, R. A. Nishimura, R. O. Bonow, B. A. Carabello, J. P. Erwin III, F. Gentile, H. Jneid, E. V. Krieger, M. Mack, C. McLeod, et al., 2020 acc/aha guideline for the management of patients with valvular heart disease: executive summary: a report of the american college of cardiology/american heart association joint committee on clinical practice guidelines, Journal of the American College of Cardiology 77 (2021) 450–500.

[3] M. J. Mack, M. B. Leon, V. H. Thourani, P. Pibarot, R. T. Hahn, P. Genereux, S. K. Kodali, S. R. Kapadia, D. J. Cohen, S. J. Pocock, et al., Transcatheter aortic-valve replacement in low-risk patients at five years, New England Journal of Medicine 389 (2023) 1949–1960.

[4] J. K. Forrest, G. M. Deeb, S. J. Yakubov, H. Gada, M. A. Mumtaz Ramlawi, T. Bajwa, P. S. Teirstein, D. Tchétché, J. Huang, et al., 4-year outcomes of patients with aortic stenosis in the evolut low risk trial, Journal of the American College of Cardiology 82 (2023) 2163–2165.

[5] J. Ternacle, S. Hecht, H. Eltchaninoff, E. Salaun, M.-A. Clavel, N. Côté, P. Pibarot, Durability of transcatheter aortic valve implantation, EuroIntervention 20 (2024) e845–e864.

[6] N. Côté, P. Pibarot, M.-A. Clavel, Incidence, risk factors, clinical impact, and management of bioprosthesis structural valve degeneration, Curr. Opin. Cardiol. 32 (2017) 123–129.

[7] A. E. Kostyunin, A. E. Yuzhalin, M. A. Rezvova, E. A. Ovcharenko, T. V. Glushkova, A. G. Kutikhin, Degeneration of bioprosthetic heart valves: update 2020, Journal of the American Heart Association 9 (2020) e018506.

[8] S. J. Bozso, J. J. Kang, R. Basu, B. Adam, J. R. Dyck, G. Y. Oudit, M. C. Moon, D. H. Freed, J. Nagendran, J. Nagendran, Structural valve deterioration is linked to increased immune infiltrate and chemokine expression, Journal of Cardiovascular Translational Research 14 (2021) 503–512.

[9] S. Wen, Y. Zhou, W. Y. Yim, S. Wang, L. Xu, J. Shi, W. Qiao, N. Dong, Mechanisms and drug therapies of bioprosthetic heart valve calcification, Frontiers in Pharmacology 13 (2022) 909801.

[10] T. Rheude, C. Pellegrini, S. Cassese, J. Wiebe, S. Wagner, T. Trenkwalder, H. Alvarez, P. Mayr, C. Hengstenberg, H. Schunkert, A. Kastrati, O. Husser, M. Joner, Predictors of haemodynamic structural valve deterioration following transcatheter aortic valve implantation with latest-generation balloon-expandable valves, EuroIntervention 15 (2020) 1233–1239. URL: https://doi.org/10.4244/EIJ-D-19-00710. doi:10.4244/EIJ-D-19-00710.

[11] M. Guglielmo, L. Fusini, M. Muratori, G. Tamborini, V. Mantegazza Andreini, A. Annoni, M. Babbaro, A. Baggiano, E. Conte, et al., Computed tomography predictors of structural valve degeneration in patients undergoing transcatheter aortic valve implantation with balloon-expandable prostheses, European Radiology (2022) 1–11.

[12] T. D. Tnay, D. Shell, A. Lui, Review of bioprosthetic structural valve deterioration: Patient or valve?, Journal of Cardiac Surgery 37 (2022) 5243–5253.

[13] A. Ochi, K. Cheng, B. Zhao, A. A. Hardikar, K. Negishi, Patient risk factors for bioprosthetic aortic valve degeneration: a systematic review and meta-analysis, Heart, Lung and Circulation 29 (2020) 668–678.

[14] D. Carbonaro, D. Gallo, U. Morbiducci, A. Audenino, C. Chiastra, In silico biomechanical design of the metal frame of transcatheter aortic valves: multi-objective shape and cross-sectional size optimization, Structural and Multidisciplinary Optimization 64 (2021) 1825–1842.

[15] B. Grossi, S. Barati, A. Ramella, F. Migliavacca, J. F. R. Matas, G. Dubini, N. Chakfé, F. Heim, O. Cozzi, G. Condorelli, et al., Validation evidence with experimental and clinical data to establish credibility of tavi patient-specific simulations, Computers in Biology and Medicine 182 (2024) 109159.

[16] G. Luraghi, F. Migliavacca, A. García-González, C. Chiastra, A. Rossi, B. Cao, G. Stefanini, J. F. R. Matas, On the modeling of patient-specific transcatheter aortic valve replacement: a fluid–structure interaction approach, Cardiovascular engineering and technology 10 (2019) 437–455.

[17] A. A. Basri, M. Zuber, E. I. Basri, M. S. Zakaria, A. F. Aziz, M. Tamagawa, K. A. Ahmad, Fluid structure interaction on paravalvular leakage of transcatheter aortic valve implantation related to aortic stenosis: A patient-specific case, Computational and mathematical methods in medicine 2020 (2020) 9163085.

[18] M. Bianchi, G. Marom, R. P. Ghosh, O. M. Rotman, P. Parikh, L. Gruberg, D. Bluestein, Patient-specific simulation of transcatheter aortic valve replacement: impact of deployment options on paravalvular leakage, Biomechanics and modeling in mechanobiology 18 (2019) 435–451.

[19] W. Mao, Q. Wang, S. Kodali, W. Sun, Numerical parametric study of paravalvular leak following a transcatheter aortic valve deployment into a patient-specific aortic root, Journal of biomechanical engineering 140 (2018) 101007.

[20] G. M. Bosi, C. Capelli, M. H. Cheang, N. Delahunty, M. Mullen, A. M. Taylor, S. Schievano, A validated computational framework to predict outcomes in tavi, Scientific reports 10 (2020) 1–11.

[21] G. Rocatello, N. El Faquir, G. De Santis, F. Iannaccone, J. Bosmans, O. De Backer, L. Sondergaard, P. Segers, M. De Beule, P. de Jaegere, et al., Patient-specific computer simulation to elucidate the role of contact pressure in the development of new conduction abnormalities after catheter-based implantation of a self-expanding aortic valve, Circulation: Cardiovascular Interventions 11 (2018) e005344.

[22] M. Heitkemper, H. Hatoum, A. Azimian, B. Yeats, J. Dollery, B. Whitson, G. Rushing, J. Crestanello, S. M. Lilly, L. P. Dasi, Modeling risk of coronary obstruction during transcatheter aortic valve replacement, The Journal of thoracic and cardiovascular surgery 159 (2020) 829–838.

[23] C. Martin, W. Sun, Simulation of long-term fatigue damage in bioprosthetic heart valves: effects of leaflet and stent elastic properties, Biomechanics and modeling in mechanobiology 13 (2014) 759–770.

[24] F. Sulejmani, A. Caballero, C. Martin, T. Pham, W. Sun, Evaluation of transcatheter heart valve biomaterials: computational modeling using bovine and porcine pericardium, Journal of the mechanical behavior of biomedical materials 97 (2019) 159–170.

[25] V. Stanová, R. Rieu, L. Thollon, E. Salaun, J. Rodés-Cabau, N. Côté, D. Mantovani, P. Pibarot, Leaflet mechanical stress in different designs and generations of transcatheter aortic valves: an in vitro study, Structural Heart 8 (2024) 100262.

[26] E. Tsolaki, P. Corso, R. Zboray, J. Avaro, C. Appel, M. Liebi, S. Bertazzo, P. P. Heinisch, T. Carrel, D. Obrist, et al., Multiscale multimodal characterization and simulation of structural alterations in failed bioprosthetic heart valves, Acta Biomaterialia 169 (2023) 138–154.

[27] I. Fumagalli, M. Fedele, C. Vergara, S. Ippolito, F. Nicolò, C. Antona, R. Scrofani, A. Quarteroni, et al., An image-based computational hemodynamics study of the systolic anterior motion of the mitral valve, Computers in Biology and Medicine 123 (2020) 103922.

[28] L. Crugnola, C. Vergara, L. Fusini, I. Fumagalli, G. Luraghi, A. Redaelli, G. Pontone, Computational hemodynamic indices to identify transcatheter aortic valve implantation degeneration, Computer Methods and Programs in Biomedicine 259 (2025) 108517.

[29] I. Fumagalli, R. Polidori, F. Renzi, L. Fusini, A. Quarteroni, G. Pontone, C. Vergara, Fluid-structure interaction analysis of transcatheter aortic valve implantation, International Journal for Numerical Methods in Biomedical Engineering 39 (2023) e3704.

[30] R. S. Rao, H. Maniar, A. Zajarias, Sapien valve: past, present, and future, Cardiac Interv Today 9 (2015) 35–41.

[31] D. Dvir, T. Bourguignon, C. M. Otto, R. T. Hahn, R. Rosenhek, J. G. Webb, H. Treede, M. E. Sarano, T. Feldman, H. C. Wijeysundera, et al., Standardized definition of structural valve degeneration for surgical and transcatheter bioprosthetic aortic valves, Circulation 137 (2018) 388–399.

[32] L. Antiga, M. Piccinelli, L. Botti, B. Ene-Iordache, A. Remuzzi, D. A. Steinman, An image-based modeling framework for patient-specific computational hemodynamics, Medical & biological engineering & computing 46 (2008) 1097–1112.

[33] T. Rheude, J. Blumenstein, H. Möllmann, O. Husser, Spotlight on the sapien 3 transcatheter heart valve, Medical Devices: Evidence and Research (2018) 353–360.

[34] C. Catalano, T. Turgut, O. Zahalka, N. Götzen, S. Cannata, G. Gentile, V. Agnese, C. Gandolfo, S. Pasta, On the material constitutive behavior of the aortic root in patients with transcatheter aortic valve implantation, Cardiovascular Engineering and Technology 15 (2024) 95–109.

[35] N. Holoshitz, C. J. Kavinsky, Z. M. Hijazi, The edwards sapien transcatheter heart valve for calcific aortic stenosis: a review of the valve, procedure, and current literature, Cardiology and therapy 1 (2012) 6.

[36] K. Capellini, E. Vignali, E. Costa, E. Gasparotti, M. E. Biancolini Landini, V. Positano, S. Celi, Computational fluid dynamic study for ataa hemodynamics: an integrated image-based and radial basis functions mesh morphing approach, Journal of biomechanical engineering 140 (2018).

[37] O. Rossvoll, S. Samstad, H. G. Torp, D. T. Linker, T. Skjærpe, B. A. Angelsen, L. Hatle, The velocity distribution in the aortic anulus in normal subjects: a quantitative analysis of two-dimensional doppler flow maps, Journal of the American Society of Echocardiography 4 (1991) 367–378.

[38] A. D. Caballero, S. Laín, A review on computational fluid dynamics modelling in human thoracic aorta, Cardiovascular Engineering and Technology 4 (2013) 103–130.

[39] A. Leuprecht, S. Kozerke, P. Boesiger, K. Perktold, Blood flow in the human ascending aorta: a combined mri and cfd study, Journal of engineering mathematics 47 (2003) 387–404.

[40] P. D. Stein, H. N. Sabbah, Turbulent blood flow in the ascending aorta of humans with normal and diseased aortic valves., Circulation Research 39 (1976) 58–65.

[41] E. L. Manchester, S. Pirola, M. Y. Salmasi, D. P. O’Regan, T. Athanasiou, X. Y. Xu, Analysis of turbulence effects in a patient-specific aorta with aortic valve stenosis, Cardiovascular engineering and technology 12 (2021) 438–453.

[42] F. Nicoud, H. B. Toda, O. Cabrit, S. Bose, J. Lee, Using singular values to build a subgrid-scale model for large eddy simulations, Physics of Fluids 23 (2011) 085106.

[43] M. Rieth, F. Proch, O. Stein, M. Pettit, A. Kempf, Comparison of the sigma and smagorinsky les models for grid generated turbulence and a channel flow, Computers & Fluids 99 (2014) 172–181.

[44] M. Fedele, E. Faggiano, L. Dedè, A. Quarteroni, A patient-specific aortic valve model based on moving resistive immersed implicit surfaces, Biomech. Model. Mechanobiol. 16 (2017) 1779–1803.

[45] A. Veneziani, C. Vergara, Flow rate defective boundary conditions in haemodynamics simulations, International Journal for Numerical Methods in Fluids 47 (2005) 803–816.

[46] L. Formaggia, J.-F. Gerbeau, F. Nobile, A. Quarteroni, Numerical treatment of defective boundary conditions for the navier–stokes equations, SIAM Journal on Numerical Analysis 40 (2002) 376–401.

[47] L. Bennati, Image-based numerical modeling of blood dynamics in presence of regurgitant and repaired mitral valve, Phd thesis, Università degli studi di Verona, 2024. Available at https://iris.univr.it/handle/11562/1145367.

[48] T. E. Tezduyar, Stabilized finite element formulations for incompressible flow computations, Advances in applied mechanics 28 (1991) 1–44.

[49] M. E. Moghadam, Y. Bazilevs, T.-Y. Hsia, I. E. Vignon-Clementel, A. L. Marsden, A comparison of outlet boundary treatments for prevention of backflow divergence with relevance to blood flow simulations, Computational Mechanics 48 (2011) 277–291.

[50] Y. Saad, Iterative methods for sparse linear systems, SIAM, 2003.

[51] L. Crugnola, C. Vergara, Inexact block lu preconditioners for incompressible fluids with flow rate conditions, arXiv preprint arXiv:2411.03929 (2024).

[52] P. C. Africa, lifex: A flexible, high performance library for the numerical solution of complex finite element problems, SoftwareX 20 (2022) 101252.

[53] P. C. Africa, I. Fumagalli, M. Bucelli, A. Zingaro, M. Fedele, A. Quarteroni, et al., lifex-cfd: An open-source computational fluid dynamics solver for cardiovascular applications, Computer Physics Communications 296 (2024) 109039.

[54] U. Morbiducci, V. Mazzi, M. Domanin, G. De Nisco, C. Vergara, D. A. Steinman, D. Gallo, Wall shear stress topological skeleton independently predicts long-term restenosis after carotid bifurcation endarterectomy, Annals of biomedical engineering 48 (2020) 2936–2949.

[55] L. Chen, H. Deng, H. Cui, J. Fang, Z. Zuo, J. Deng, Y. Li, X. Wang, L. Zhao, Inflammatory responses and inflammation-associated diseases in organs, Oncotarget 9 (2017) 7204.

[56] P. Eshtehardi, A. J. Brown, A. Bhargava, C. Costopoulos, O. Y. Hung, M. T. Corban, H. Hosseini, B. D. Gogas, D. P. Giddens, H. Samady, High wall shear stress and high-risk plaque: an emerging concept, The international journal of cardiovascular imaging 33 (2017) 1089–1099.

[57] A. Arzani, S. C. Shadden, Wall shear stress fixed points in cardiovascular fluid mechanics, Journal of biomechanics 73 (2018) 145–152.

[58] L. Chen, H. Qu, B. Liu, B.-C. Chen, Z. Yang, D.-Z. Shi, Y. Zhang, Low or oscillatory shear stress and endothelial permeability in atherosclerosis, Frontiers in Physiology 15 (2024) 1432719.

[59] Y. Kang, H. Y. Hwang, S. H. Sohn, J. W. Choi, K. H. Kim, K.-B. Kim, Comparative analysis of structural valve deterioration after bioprosthetic tricuspid valve replacement: bovine pericardial versus porcine valves, Artificial Organs 45 (2021) 911–918.

[60] P. Di Achille, G. Tellides, C. Figueroa, J. Humphrey, A haemodynamic predictor of intraluminal thrombus formation in abdominal aortic aneurysms, Proceedings of the Royal Society A: Mathematical, Physical and Engineering Sciences 470 (2014) 20140163.

[61] H. Wang, D. Balzani, V. Vedula, K. Uhlmann, F. Varnik, On the potential self-amplification of aneurysms due to tissue degradation and blood flow revealed from fsi simulations, Frontiers in Physiology 12 (2021) 785780.

[62] C. Trenti, M. Ziegler, N. BjarnegÃ, T. Ebbers, M. Lindenberger, P. Dyverfeldt, et al., Wall shear stress and relative residence time as potential risk factors for abdominal aortic aneurysms in males: a 4d flow cardiovascular magnetic resonance caseâ€”control study, Journal of Cardiovascular Magnetic Resonance 24 (2022) 18.

[63] E. Y. Lui, A. H. Steinman, R. S. Cobbold, K. W. Johnston, Human factors as a source of error in peak doppler velocity measurement, Journal of vascular surgery 42 (2005) 972–e1.

[64] J. Niederberger, H. Schima, G. Maurer, H. Baumgartner, Importance of pressure recovery for the assessment of aortic stenosis by doppler ultrasound: role of aortic size, aortic valve area, and direction of the stenotic jet in vitro, Circulation 94 (1996) 1934–1940.

[65] R. F. Siddiqui, J. R. Abraham, J. Butany, Bioprosthetic heart valves: modes of failure, Histopathology 55 (2009) 135–144.

[66] S. Madhavan, E. M. C. Kemmerling, The effect of inlet and outlet boundary conditions in image-based cfd modeling of aortic flow, Biomedical engineering online 17 (2018) 66.

[67] A. Quarteroni, A. Manzoni, C. Vergara, The cardiovascular system: mathematical modelling, numerical algorithms and clinical applications, Acta Numerica 26 (2017) 365–590.

[68] F. L. Abel, Influence of aortic compliance on coronary blood flow., Circulatory shock 12 (1984) 265–276.

[69] J. Li, W. Yan, W. Wang, S. Wang, L. Wei, Comparison of balloon-expandable valve and self-expandable valve in transcatheter aortic valve replacement: A patient-specific numerical study, Journal of Biomechanical Engineering 144 (2022) 104501.

[70] F. Duca, D. Bissacco, L. Crugnola, C. Faitini, M. Domanin, F. Migliavacca, S. Trimarchi, C. Vergara, Computational analysis to assess hemodynamic forces in descending thoracic aortic aneurysms, The Journal of Physiology (2025).

